# Revealing the Assembly of Filamentous Proteins with Scanning Transmission Electron Microscopy

**DOI:** 10.1101/697649

**Authors:** Cristina Martinez-Torres, Federica Burla, Celine Alkemade, Gijsje H. Koenderink

## Abstract

Filamentous proteins are responsible for the superior mechanical strength of our cells and tissues. The remarkable mechanical properties of protein filaments are tied to their complex molecular packing structure. However, since these filaments have widths of several to tens of nanometers, it has remained challenging to quantitatively probe their molecular mass density and three-dimensional packing order. Scanning transmission electron microscopy (STEM) is a powerful tool to perform simultaneous mass and morphology measurements on filamentous proteins at high resolution, but its applicability has been greatly limited by the lack of automated image processing methods. Here, we demonstrate a semi-automated tracking algorithm that is capable of analyzing the molecular packing density of intra- and extracellular protein filaments over a broad mass range from STEM images. We prove the wide applicability of the technique by analyzing the mass densities of two cytoskeletal proteins (actin and microtubules) and of the main protein in the extracellular matrix, collagen. The high-throughput and spatial resolution of our approach allow us to quantify the internal packing of these filaments and their polymorphism by correlating mass and morphology information. Moreover, we are able to identify periodic mass variations in collagen fibrils that reveal details of their axially ordered longitudinal self-assembly. STEM-based mass mapping coupled with our tracking algorithm is therefore a powerful technique in the characterization of a wide range of biological and synthetic filaments.

## MAIN TEXT

The main structural components of cells and tissues are scaffolds made of proteins that self-assemble into filaments. Cells are sustained by cytoskeletal filaments, whilst tissues are supported by an extracellular matrix composed predominantly of collagen fibrils. These protein scaffolds have unique material properties, combining a superior mechanical resistance to large deformations with the ability to dynamically adapt, grow and repair.^1–3^ There is growing evidence that the molecular packing structure of protein filaments is an important determinant of these unique material properties. Both cytoskeletal and extracellular filaments are supramolecular assemblies with a highly organized molecular structure dictated by specific noncovalent interactions between the constituent proteins. For a quantitative understanding of the relation between the mechanics and structure of these biopolymers, we need to be able to quantitatively characterize their internal molecular packing arrangement. This task is challenging as biopolymers are thin structures (with widths of 5 – 25 nm for cytoskeletal filaments and 10 – 500 nm for extracellular filaments) with small masses (1 kDa – 1 MDa). One powerful tool to spatially resolve the mass density of supramolecular protein assemblies is scanning transmission electron microscopy. In this technique, a converging beam of electrons scans across a thin specimen and by collecting only those electrons that are scattered at very high angle, the so-called High Angle Angular Dark-Field Mode (HAADF) (Fig. 1A), the resulting image intensity is directly proportional to the mass of the specimen.^4, 5^ The proportionality constant necessary to satisfy this relation can be obtained either by an extensive calibration of the electron microscope,^6,7^ or by imaging the protein of interest together with a reference specimen that serves as an internal mass calibration.^5,8^

**Figure 1.**
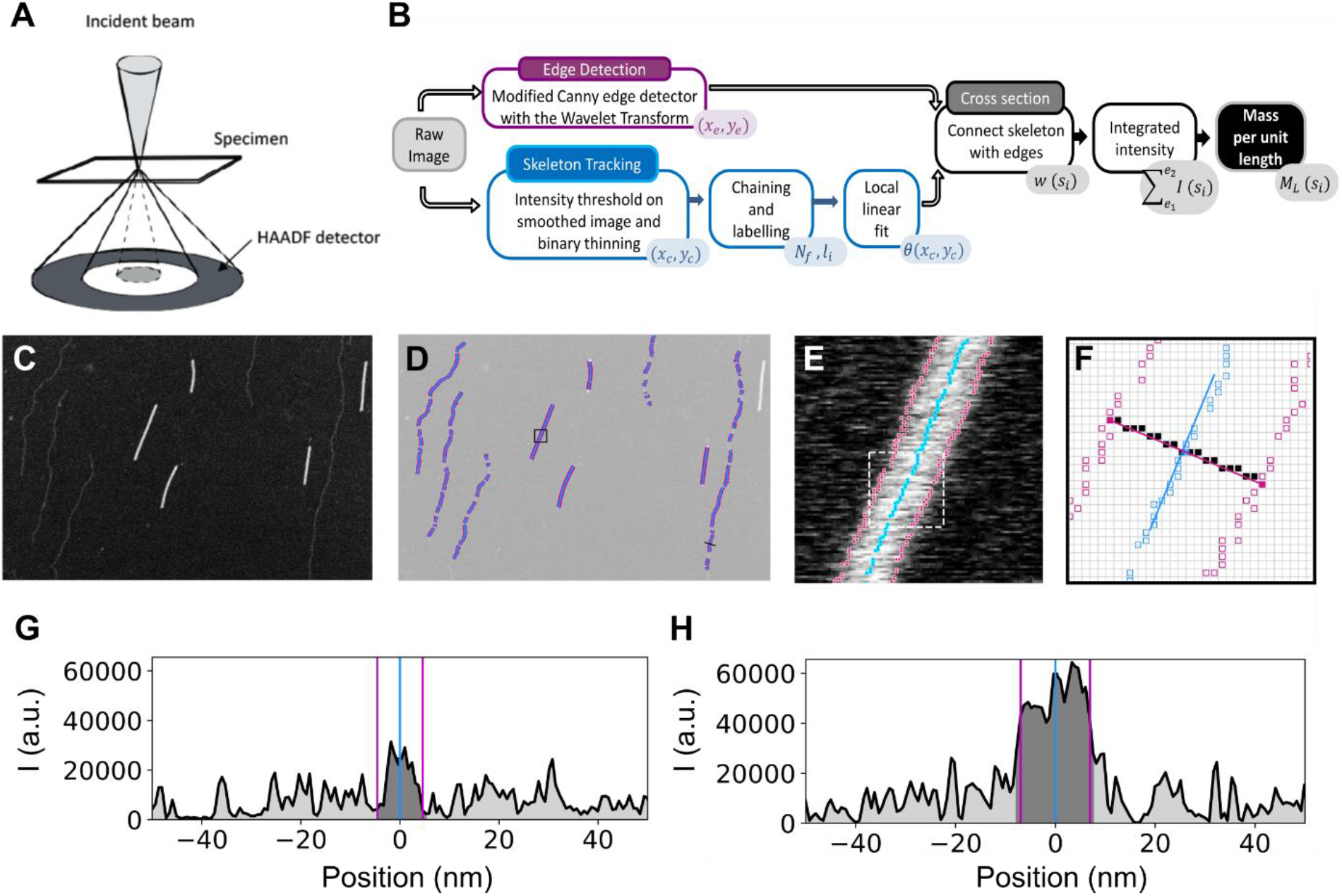
Overview of the method for filament tracking and mass-per-length (*M*_*L*_) computation. (A) Scheme of the HAAFD mode, where the incident electrons are scattered by the specimen and collected at high angles with an angular detector. (B) Algorithm workflow showing the steps and the corresponding output (color-filled text boxes in the lower right corners). (C) Unprocessed image of TMV rods (bright short rods) and Fd bacteriophages (dim long filaments) serving as input in this example. (D) Tracked edges (magenta) and skeleton (blue) points, used to compute the *M*_*L*_. Only the filament segments with a clean background and without intensity saturation are retained. (E) Image inside the black square shown in (D), with the tracked points overlaid. (F) Tracked edges and skeleton points inside the dashed white square shown in (E). The blue line shows the linear fit to the skeleton point in the center while the corresponding cross section is shown in magenta. The solid black squares represent the pixels in the filament cross section. (G) Intensity profile of the cross-section of the TMV rod in (F). The vertical lines show the detected edges and skeleton. Dark and light grey represent the integrated intensity for the filament and the background, respectively. (H) Same as (G) for an Fd filament indicated with a black line in (D).

Although mass mapping with STEM has already been performed on multiple different systems,^9–13^ its applicability and the information content have been greatly limited by the approach used for data analysis. To obtain the mass per unit length of the filaments, typically a manual image analysis is performed: a box is drawn around the filament of interest, and the integrated intensity is computed inside this region, taking into account a correction for the background intensity surrounding the filament. This approach has several drawbacks. First, there is the limited statistics that can be reasonably reached by manually selecting the coordinates and/or width of the bounding box. Second, manual analysis requires user input in defining the bounding box, which could potentially introduce a considerable human bias in the results, particularly if the width of the filament is one of the variables to be analyzed. Finally, the discretization of the points along the filament axis could mask interesting features about spatial modulations in protein density along or across the filament axis. Many natural proteins form filaments that are highly ordered due to precise axially staggered self-assembly. The best-known example is the extracellular matrix protein collagen, a long rod-shaped molecule that assembles in a quarter-staggered manner to form filaments with a 67 nm axial periodicity referred to as the D-period.^11,14^

Here, we report a semi-automated algorithm to retrieve the mass per length of filamentous structures and obtain high throughput information on the correlation between filament mass and morphology. Our method automatically tracks the filament skeleton and its borders via a modified canny edge detection with the wavelet transform,^15–17^ resulting in a robust tracking even in conditions of low image contrast for proteins of low density. The principle of this analysis is to smooth the image and compute its gradient, by convolving the image with an appropriate filter, such as the first derivative of a Gaussian function. This approach allows us to retrieve the distribution of mass per length and the widths for all fibers within a given image. Moreover, it enables us to quantify detailed structural features of fiber and network assembly, such as fiber internal packing, axial packing periodicity, fiber twisting and branching. The method we propose therefore opens up the way to quantitatively dissect the assembly of complex filamentous structures, from natural to synthetic fiber-forming systems.

## Results and Discussion

The workflow of our algorithm is summarized in Figure 1B. The starting point is the unprocessed HAADF image (Fig. 1C), where the intensity is minimal for the background (black) and maximal for the pixels containing the specimen that scatter the electrons (white). The method follows two independent tracking processes, one to detect the filament edges, and the other one to detect the filament main axis, or skeleton (Fig. 1D-E). The skeleton is detected by thinning a binary mask obtained by thresholding the intensity of the image (smoothed by a Gaussian filter). Next, neighboring points are connected into chains by a 8-neighbour rule and each chain is labeled with a different number to identify connected points. Short chains may be identified as noise and discarded at this point. The final step in tracking the filament skeleton performs a local linear fit around each point (Fig. 1F), in order to obtain the local orientation of the filament and to retrieve transverse intensity profiles all along the axis of the filament. The fit range can be modified depending on the persistence length of the filament. In the edge tracking algorithm, we use the first derivative of a Gaussian as a kernel. When using this function in the wavelet transform, the result is a smoothed version of the 2D gradient of the image.^17^ Moreover, the transform gives an argument and a modulus for each point, that can be used as input for a Canny detection algorithm. At each image coordinate, a point is considered an edge if the wavelet transform modulus is a local maximum when compared to its neighboring pixels. The comparison is made with those pixels that follow the argument of the wavelet transform at that point. To identify ‘true’ edges, the points are chained together and a double hysteresis algorithm is applied to connect weak with strong edges.^15^ Once the filament edges have been detected, they can be assigned to a point in the filament axis (skeleton), by looking for them in the cross-section profile (Fig. 1F). The integrated density is then computed for the identified filament region, subtracting the intensity of the background on each side of the filament to correct for any scattering from the background and the support film (Fig. 1G-H). If the constant of proportionality between the intensity and mass per length (*M*_*L*_) is known, the integrated intensity can be converted to *M*_*L*_ values. In this study, we use tobacco mosaic virus (TMV) rods for mass calibration, since these have a constant and well-known mass per length ratio.^18^ However, depending on the morphology and mass range of the filaments to be studied, other well-calibrated structures can be used as reference, for example Fd bacteriophages,^19^ which appear as dim long filaments in Fig. 1C and which likewise have a constant and well-known mass per length ratio (Fig. S1).

### Method validation on virus rods and cytoskeletal filaments

The cell owes its shape and mechanical properties to a filamentous scaffold known as the cytoskeleton. Two of the structural proteins forming the backbone of the cytoskeleton, actin and microtubules, are known to assemble into relatively simple and monodisperse structures. Therefore, we decided to use these proteins as test structures to validate our method. Actin filaments are helical polymers composed of two linear strands of globular actin subunits (Fig. 2A, inset). With a width of only 8 nm and *M*_*L*_ of 16 kDa/nm,^9,20–21^ an actin filament is one of the smallest structures we can measure given the accuracy of STEM mass mapping: in general an image resolution of 2-4 nm can be achieved depending on the sample preparation, while the mass range that can be detected is typically in the range of a few 10 kDa up to 100 MDa.^4,22^ Figure 2A shows a HAADF image of an actin filament deposited on an EM grid covered with a 3 nm thick carbon film. Although it is possible to reach a higher resolution by increasing the magnification of the microscope, it is important to keep the electron dose per nm^2^ to a minimum to prevent mass loss of the protein,^5^ and thus, an incorrect mass quantification. We optimized the imaging settings to enhance contrast, while maintaining the intensity of the reference TMV rods (lower right corner in Fig. 2A) just below saturation. Our tracking method recovers an homogeneous distribution of *M*_*L*_ values, with <*M*_*L*_> = 17.22 ± 6.91 kDa/nm and an average width <w> = 15.77 ± 1.82 nm. The width of the filaments is larger than expected, which is likely explained by two factors. On the one hand, the spatial resolution is limited since 1 pixel typically corresponds to ~ 1 nm and the edge tracking method has a maximum accuracy of 2 nm in determining the width of a filament (± 1 pixel on each border). We can thus expect an error on the order of 2-4 nm depending on the image resolution. On the other hand, the sample preparation can also change the width of the fibrils compared to the hydrated diameter in solution due to surface adsorption and drying. Since we avoided chemical fixation in order not to change the *M*_*L*_, the actin filaments likely flatten upon surface adsorption. We next tested our method on microtubules, which form hollow sheets made of 12-15 linear protofilaments^23^ (see inset of Fig. 2B). Our tracking method recovers an <*M*_*L*_ > = 178.7 ± 29.9 kDa/nm and a mean width <*w*> = 32.8 ± 3.76 nm (Fig. 2C), matching the values of native microtubules.^24^ Microtubules exhibit less filament broadening than actin filaments, perhaps due to their larger rigidity. However, they exhibit a broader spread in *M*_*L*_ values, consistent with the known variability in protofilament number.^23,25^ Given our sample preparation (see Methods), we can expect microtubules with predominantly 13 and 14 protofilaments, with a respective *M*_*L*_ of 181.5 kDa/nm and 195.4 kDa/nm, in agreement with our results. In addition, the microtubules we obtained have ends with a reduced *M*_*L*_ (Fig. 2B), likely because we did not chemically stabilize them.

**Figure 2.**
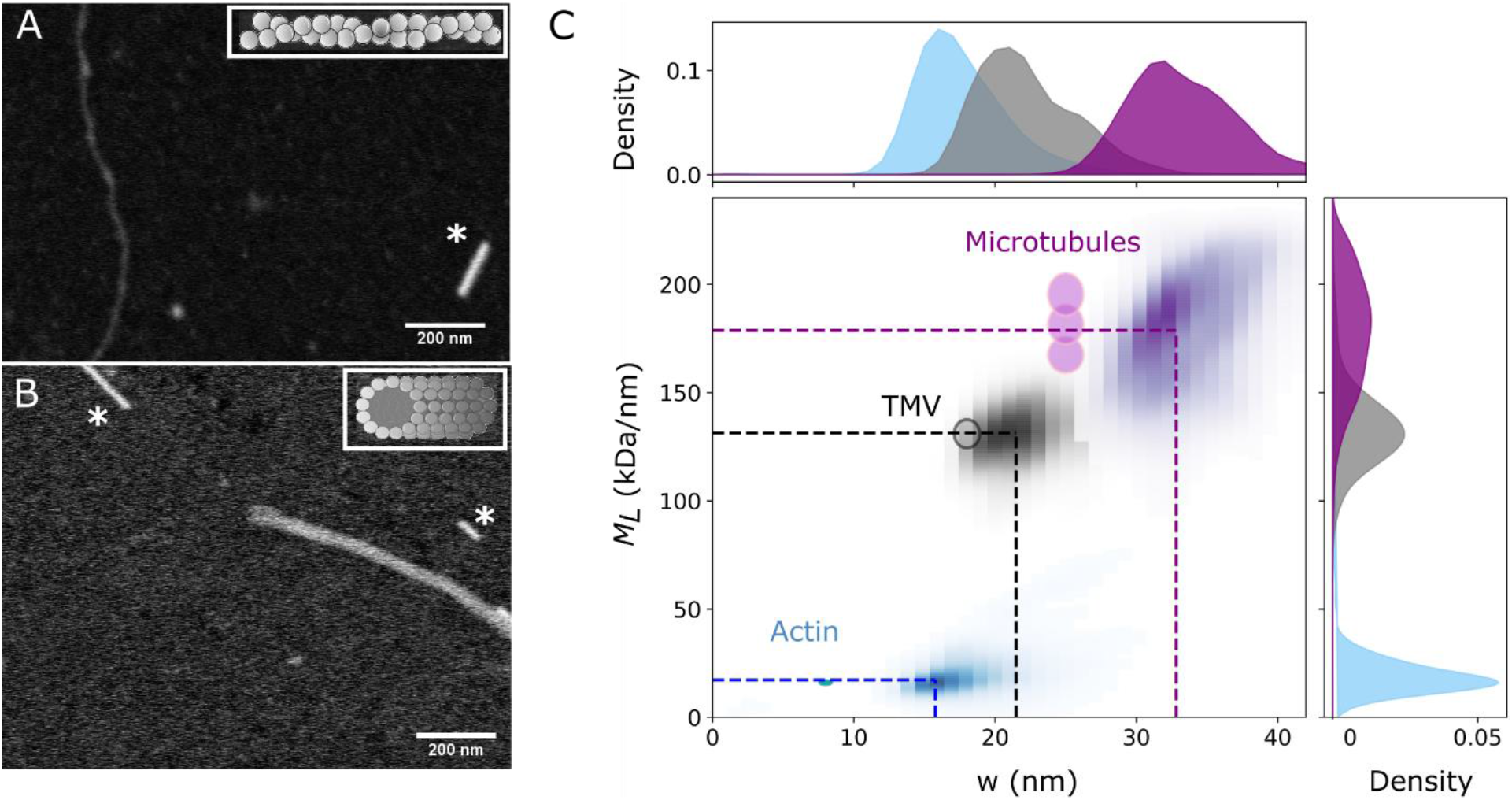
Mass mapping of cytoskeletal proteins. Unprocessed images of (A) an actin filament and (B) a microtubule. The short bright rods marked by an asterisk in both images are TMV rods used as reference. (C) Kernel density plot of the filament mass per length (*M*_*L*_) and width (w) for data collected from multiple images of microtubules (purple), TMV (black) and actin (blue). The dotted lines show the mean fitted values, and the circles indicate the expected values from literature with a ± 10 % spread. For microtubules, the three circles correspond to structures with 12, 13, and 14 protofilaments. *n* = 8150 (actin), 2986 (microtubules), 5450 (TMV) segments.

### Structural analysis of more complex fibril-forming systems

The protein mass of filamentous structures is an essential parameter indicating the number of subunits present. Above we considered actin and microtubules, both proteins with a relatively simple and well-defined assembly process. In this case the filaments form a homogeneous population and the average *M*_*L*_ and width values provide a sufficient description. However, many proteins of the extracellular matrix, like fibrin and collagen, form more complex and polymorphic supramolecular structures, and data averaging would obscure polymorphism within and between filaments. Since STEM provides concurrent information on the mass and shape of individual filaments, we propose that it is an attractive tool to study proteins that assemble into hierarchical structures. We demonstrate the potential of our filament tracking method in combination with STEM mass mapping by analyzing the assembly of collagen fibrils.

Collagen is the main structural element of the extracellular matrix of connective tissues, important for force transmission and load-bearing.^26^ The morphology of the fibrils and the architecture of collagen fibril networks vary widely among tissues, providing tissue-specific mechanical functions. Reconstitution studies have shown that collagen assembly is regulated by many environmental factors, such as temperature, pH, and ionic strength.^11,27,28^ A few studies have shown that STEM can be a useful tool to decipher the molecular pathway of collagen assembly^11^ but a detailed analysis has been lacking due to limitations in image processing. To test the applicability of STEM, we polymerize collagen fibrils from bovine dermal collagen I and adsorb them to an EM grid. As shown in Fig. 3A, the resulting fibrils are highly variable in width. The corresponding histograms of *M*_*L*_ (Fig. 3D) and width (*w*) (Fig. 3E) show several peaks, indicative of multiple populations. Note that the high output of our method uniquely reveals this polymorphic behavior. In the image analyzed (Fig. 3A), we recover the information from over 5000 transverse fibril segments. Segments where the background or the fibril were contaminated by salts or other unwanted structures, were manually filtered out from the dataset based on visual inspection.

**Figure 3.**
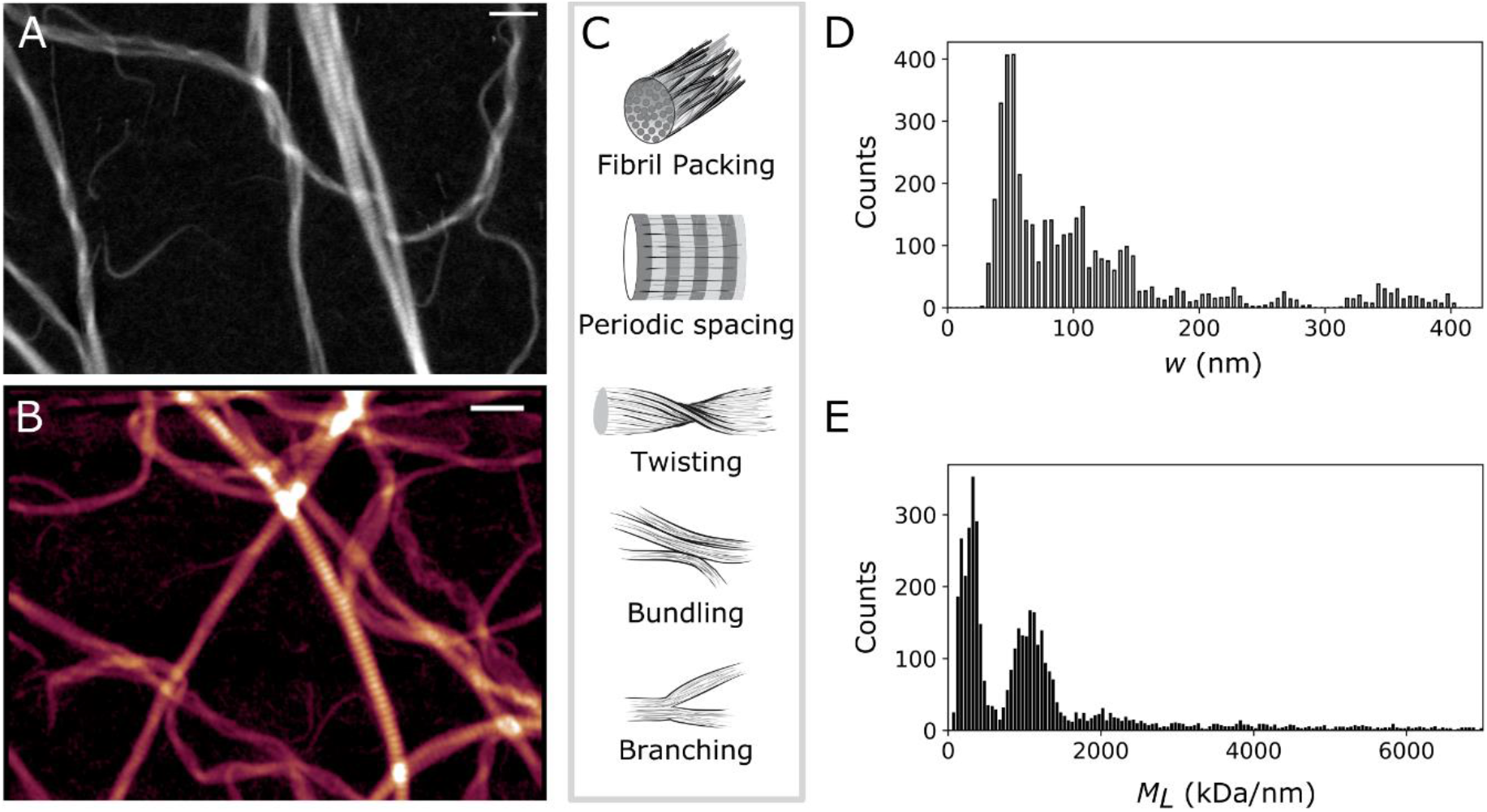
Mass mapping of bovine type I collagen fibrils. (A-B) Images of reconstituted atelocollagen fibrils adsorbed on an electron microscope grid. Images of the same sample were acquired using (A) HAADF mode with STEM, and (B) AFM contact mode in air (height color coded from 0 to 55 nm). Scale bar 500 nm. (C) Sketch of the different morphological features encountered in collagen fibrils, as seen in (A-B). (D-E) Distribution of the fibril width *w* (D) and *M*_*L*_ values (E) for the collagen fibrils in the image shown in (A). n = 5000 segments.

The EM image reveals several morphological features known to be characteristic of collagen fibrils (Fig. 3C): the fibrils are branched, bundled and twisted at large (micron) scales, and they have an axially periodic packing structure at the nanoscale.^11,14,29^ The packing periodicity is clearly visible from the regular bands with alternating regions of high and low intensity (*i.e.* mass). Furthermore, the fibrils have tapered ends, consistent with prior observations both in tissue and reconstituted collagen.^11^ The packing periodicity is often studied with atomic force microscopy^14^ (AFM) which can provide morphological, but not mass information. When we image our EM sample by AFM (Fig. 3B), we indeed observe the same qualitative features in the fibril topography as in the HAADF images of the fibril mass density (Fig. S2).

### Lateral assembly and fibril packing

The basic unit of a collagen fibril is the tropocollagen molecule, which consists of an uninterrupted triple helix of nearly 300 nm in length and 1.5 nm in diameter.^30^ In addition, the molecule may contain two short extra telopeptides, N- and C-terminal domains of 11 to 26 residues, which account for only ~2% of the molecule but which are critical in the assembly of the fibrils. The absence of these telopeptides results in a loss of diameter uniformity and changes in the fibril assembly pathway.^26,31^ Fibrils reconstituted from purified collagen usually exhibit high axial periodicity, but their degree of lateral packing order is still elusive. X-ray diffraction studies in tissue collagen indicate an almost crystalline molecular packing,^27,29,32^ but deviations from this model are often observed. The differences have been attributed to fibril curvature and twisting or to the coexistence of ordered and less-ordered domains along the fibril axis.^33,34^ However, all of these explanations remain speculative, and are mostly supported by theoretical models of molecular packing, without clear experimental evidence.

To address the question of the degree of packing density in collagen fibrils, we have studied fibrils with the presence (telo-) or absence (atelo- collagen) of the telopeptides. Rather than collapsing all the data into simple *M*_*L*_ and *w* distributions (as done in Fig. 3) we now focus on the evolution of *M*_*L*_ with the fibril diameter (*w*). Figure 4B shows the resulting curve for telocollagen (in blue) and atelocollagen (in red), where all the data (7 images with 50 fibrils for telo-, 14 images with 85 fibrils for atelo- collagen) has been pooled together, binned and averaged over a width window of 5 nm, which is the lowest imaging resolution we have in our dataset. We found that, for thick fibrils (w > 100 nm), atelocollagen fibrils have considerably lower values of *M*_*L*_ compared to telocollagen, indicating a looser molecular packing. Since the *M*_*L*_ and *w* of a single tropocollagen molecule is known (*m*_*o*_ and *w*_*o*_ respectively), we can estimate the molecular packing density according to:

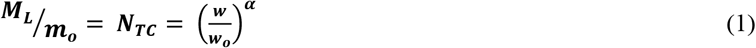

where *N*_*TC*_ is the number of tropocollagen molecules, and α is the power law describing the fibril packing density that can adopt values in the range of 1 ≤ α ≤ 2, with α = 1 (dashed gray line in Fig. 4B) corresponding to a quasi-bidimensional or flat structure, and α = 2 (dashed black line in Fig. 4B) when the molecules uniformly occupy the interior of a cylindrical fibril. For intermediate values, 1 < α < 2, we encounter a fractal structure, where the molecules are not as tightly packed. It is important to keep in mind that with STEM, as with any other technique where the biopolymer is interacting with a surface, some flattening of the fibrils can occur due to surface adsorption, which will also impact the packing density that we measure. Since we are mostly interested in the comparison between atelo- and telocollagen under the same conditions, the relative change of α values is still a meaningful parameter that is linked to the packing density in the fibrils. Using Eq. (1) to fit the data, we obtain α = 1.687 +− 0.003 for telocollagen, and α = 1.525 +− 0.003 for atelocollagen. This small difference in α translates in a large reduction in the fibril *M*_*L*_ of up to 50% for the thicker fibrils (w ~ 400 nm) when the telopeptides are absent. The difference is particularly noticeable for widths larger than ~100 nm (Fig. S3). This could signify that telocollagen fibrils have a similar packing density in the core as atelocollagen fibrils, but a tighter packing in the periphery. Another interesting observation is that we observe two packing regimes for the telocollagen fibrils: the *M*_*L*_ monotonically increases with width until ~400 nm, and thereafter increases more slowly. This threshold coincides with a clear change in morphology, from single fibrils for widths below ~400 nm to bundles above this threshold (grey-colored zone in Fig. 4B).

**Figure 4.**
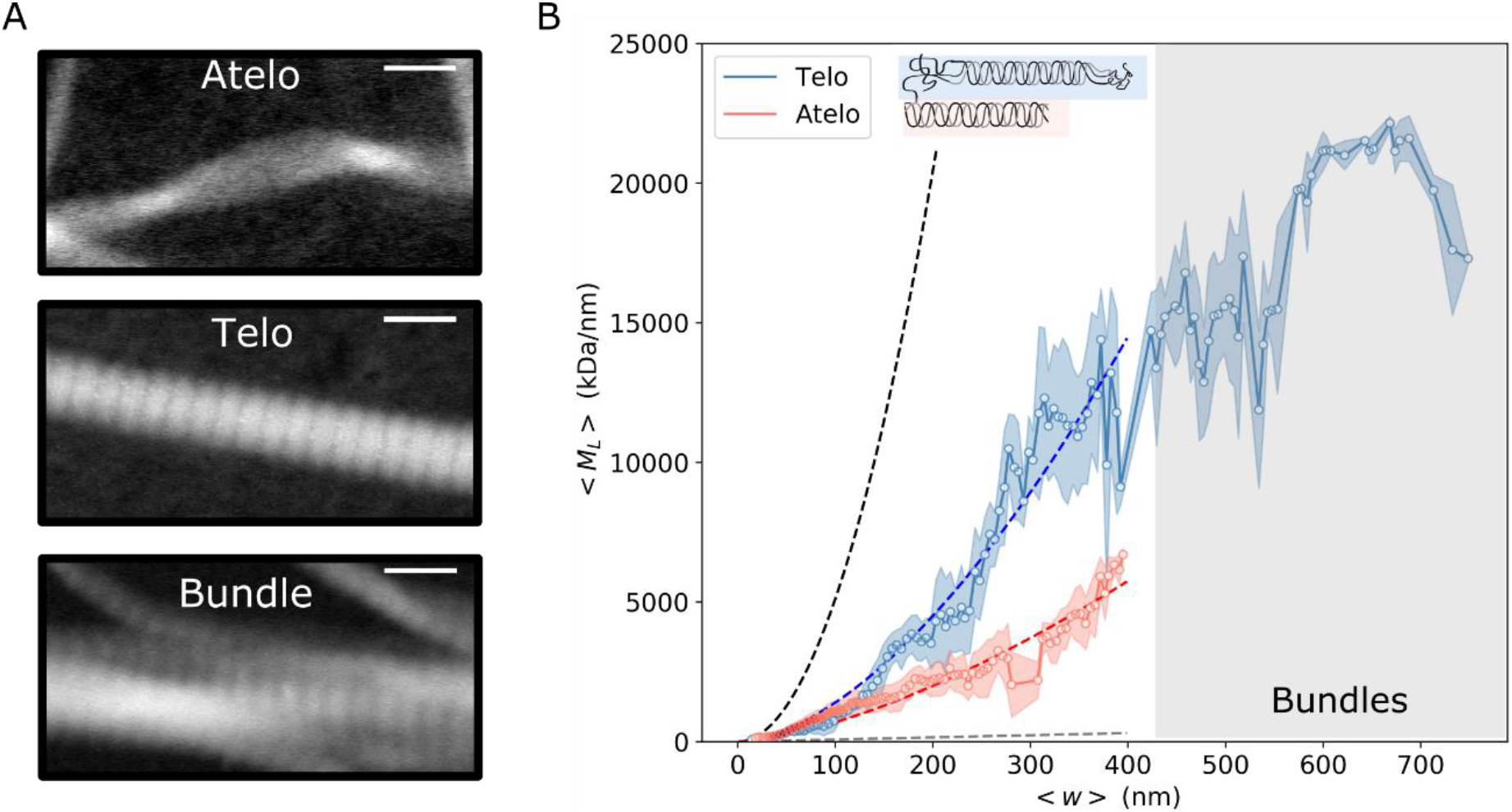
Collagen fibril packing. (A) HAADF images of representative fibrils from atelo- and telo-collagen. The bottom row shows an example of a bundle where multiple telocollagen fibrils join. Scale bar 200 nm. (B) Average *M*_*L*_ values as a function of average fibril width, *w*, for telo- (blue) and atelo- (red) bovine collagen type I. The dotted lines show the expected *M*_*L*_ values from Eq. (1) for α = 2 (black) and α = 1 (gray), and the fits with α = 1.687 and α = 1.525 for telo- (blue) and atelo-(red) collagen, respectively. Fibers in the grey-colored region are all bundles. The error bars represent the standard deviation. *n* = 15695 (telo-) and *n* = 41730 (atelo-collagen) segments.

### Longitudinal assembly and spatial features

A characteristic feature of the collagen fibrils is the so-called D-pattern or D-spacing, a periodic banded pattern appearing along the fibril axis. This axial periodicity is the result of the tropocollagen molecules aligning laterally in a quarter-staggered manner (Fig. 5A), but details of the assembly process are still being studied with the aid of different theoretical models.^14,27,31^ It has been proposed that the lateral molecular packing differs between the regions of the D-spacing, resulting in alternating regions of high packing density where the molecules overlap, and low packing density in the gap regions.^27^ To test this hypothesis, we extracted the *M*_*L*_(*w*) values for the gap and overlap regions in a collagen fibril showing a clear D-pattern (Fig. 5B). Starting from the semi-automated tracked points, we manually selected those sections that by visual inspection corresponded to either gap or overlap regions, giving a total of *n* = 547 and *n* = 733 cross-sections for ~80 gap and overlap regions, respectively. Figure 5C shows the resulting curves for the *M*_*L*_ values averaged over 5 nm width bins. Fitting the data with Eq. (1), we found α = 1.67 +− 0.002 for both the gap and overlap regions, suggesting that the alternating regions differ in their *M*_L_ but not in packing density.

**Figure 5.**
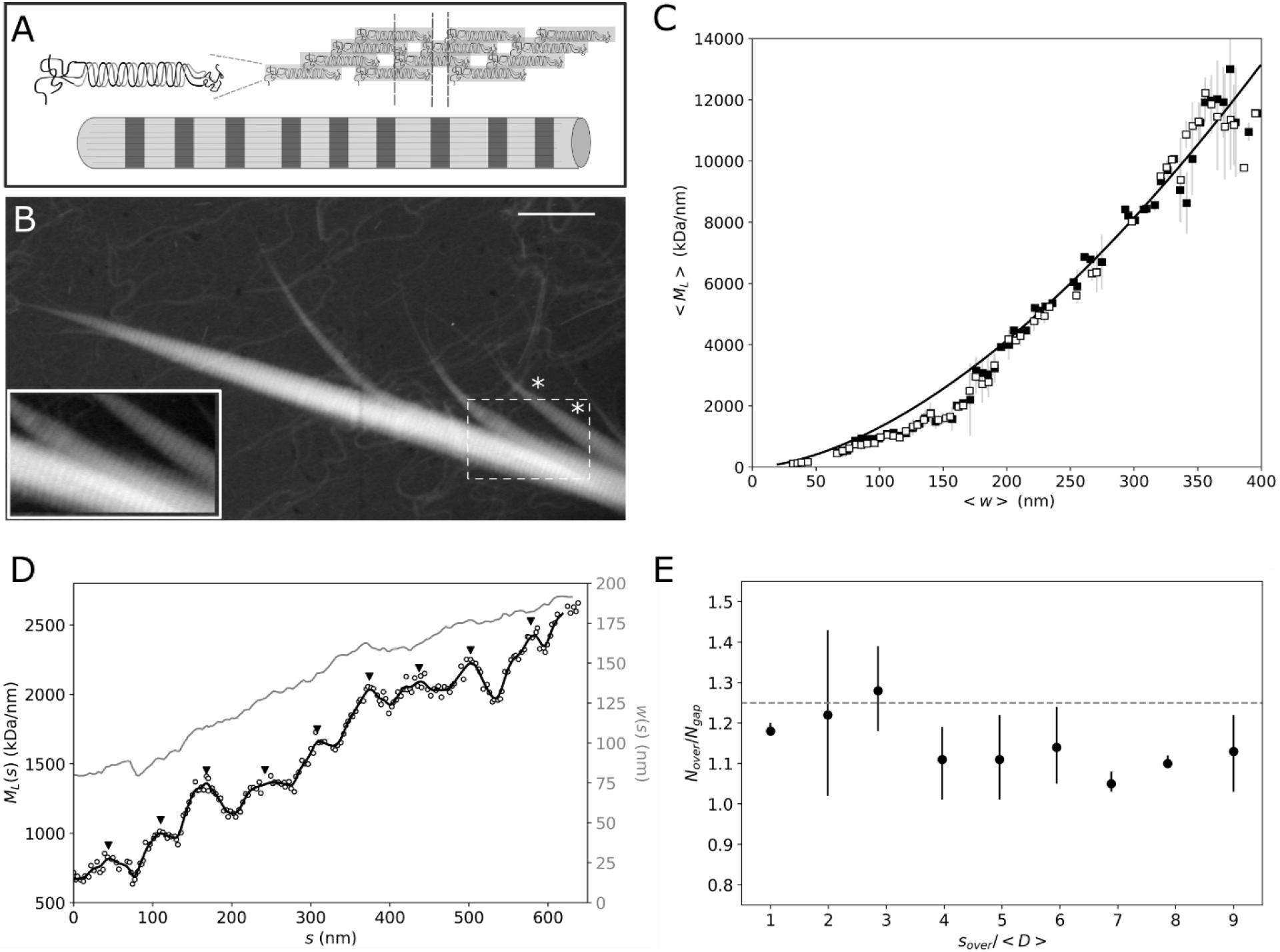
Mapping axial order in a telocollagen fibril. (A) Scheme of the quarter-staggered lateral assembly of collagen, resulting in a periodic pattern of overlap and gap regions. (B) STEM image of a single fibril from a telocollagen network. The inset is a magnification of the area in the white dashed square, where the contrast has been optimized to show the D-spacing pattern. Scale bar 1 μm. (C) Average *M*_*L*_ values as a function of *w* for the overlap (black squares) and gap (white squares) regions in the image shown in (B). The solid black line shows the fit to Eq. (1) with α = 1.67. (D) Axial mass profile for the fibril indicated with an asterisk in (B). The white circles show the raw *M*_*L*_ data, and the black line the smoothed data. The corresponding width profile is shown in gray. The black upside triangles indicate the positions of the overlap regions *s*_*over*_ (E) Ratio between the number of tropocollagen molecules in the overlap (*N*_*over*_) and gap (*N*_*gap*_) regions of the profile shown in (D). The x-axis shows the axial position of the overlap region (*s*_*over*_), normalized by the mean D-spacing of the fibril, <*D*> = 66.7 ± 5.4 nm

Our analysis also allows us to spatially map variations in *M*_*L*_ and width along the fibril axis. Figure 5D shows the *M*_*L*_ (black) and *w* (gray) profiles as a function of the fibril axis *s*, for the fibril segment enclosed by the asterisks in Fig. 5B. Both the width and mass progressively increase as we move from the pointed end to the fiber middle. This behavior is reminiscent of tapered ends known as α-tips and β-tips that have been identified for fibrils assembling from procollagen. However, those tapered ends were reported to grow by only 17 and 113 molecules/D-period, respectively,^35^ whereas for the fibril shown in Fig. 5D we find an increment of 248 ± 12 molecules/D-period, for the first 4 D-periods. This large difference in taper, together with the morphological appearance in the STEM image, suggests that the tapered ends we observe here are due to fibril splitting as a consequence of the sample preparation for EM imaging (see Methods) instead of a specific fibril growth mechanism.^35^ The spatial map of the *M*_*L*_ variation along the fibril furthermore enables us to infer a value for the average D-spacing of <*D*> = *66.7* ± *5.4* nm, consistent with prior reports based on negative stain EM and AFM.^14,26,31^ The *M*_*L*_(*s*) profile shown in Fig. 5D furthermore enables us to directly check the quarter-stagger assembly model (sketched in Fig. 5A) by measuring the ratio in *M*_*L*_ for the overlap and gap regions. We take for the positions of the overlap region, *s*_*over*_, each of the local maxima observed, and the number of tropocollagens in the overlap, *N*_*over*_, as the value of *M*_*L*_ (*s*_*over*_)/ *m*_*o*_. For the gap regions, *N*_*gap*_, we take the mean value from the local minima before and after *s*_*over*_. Figure 5E shows the ratio *N*_*over*_/*N*_*gap*_, as a function of D-period. In all the overlap regions, we get a greater number of molecules compared to the gaps, regardless of the fibril width and/or distance from the tip. In this particular case, there is a mean ratio of *N*_*over*_/*N*_*gap*_ = 1.15 +− 0.006, consistent with the theoretical ratio of 1.2 proposed in Ref.^27^

So far we have paid attention exclusively to the two main outputs of our tracking method for STEM mass mapping: the filament *M*_*L*_ and width. However, we can gain additional information by correlating this information with secondary variables involved in the processing of the images. One of the fibril features that can be studied is the intensity, which is dependent, among others, on fibril twisting. Figure 6A shows the image of a collagen fibril that appears to be twisted at the points indicated by the white asterisks, where the intensity is higher and the fibril is thinner (Fig. 6B-C). To make sure these variations in intensity are due to twist and not to differences in protein mass, we can correlate the mass profile *M*_*L*_(s) with the intensity profile *I*(*s*) (Fig. 6C). In this example the twisting points do not correlate with variations in fibril mass, suggesting that the *I*(*s*) variations are indeed due to twisting. This phenomenon is reminiscent of prior reports of twisting in collagen fibrils,^14^ although it is unclear to what extent the interaction of the fibril with the surface plays a role here.

**Figure 6.**
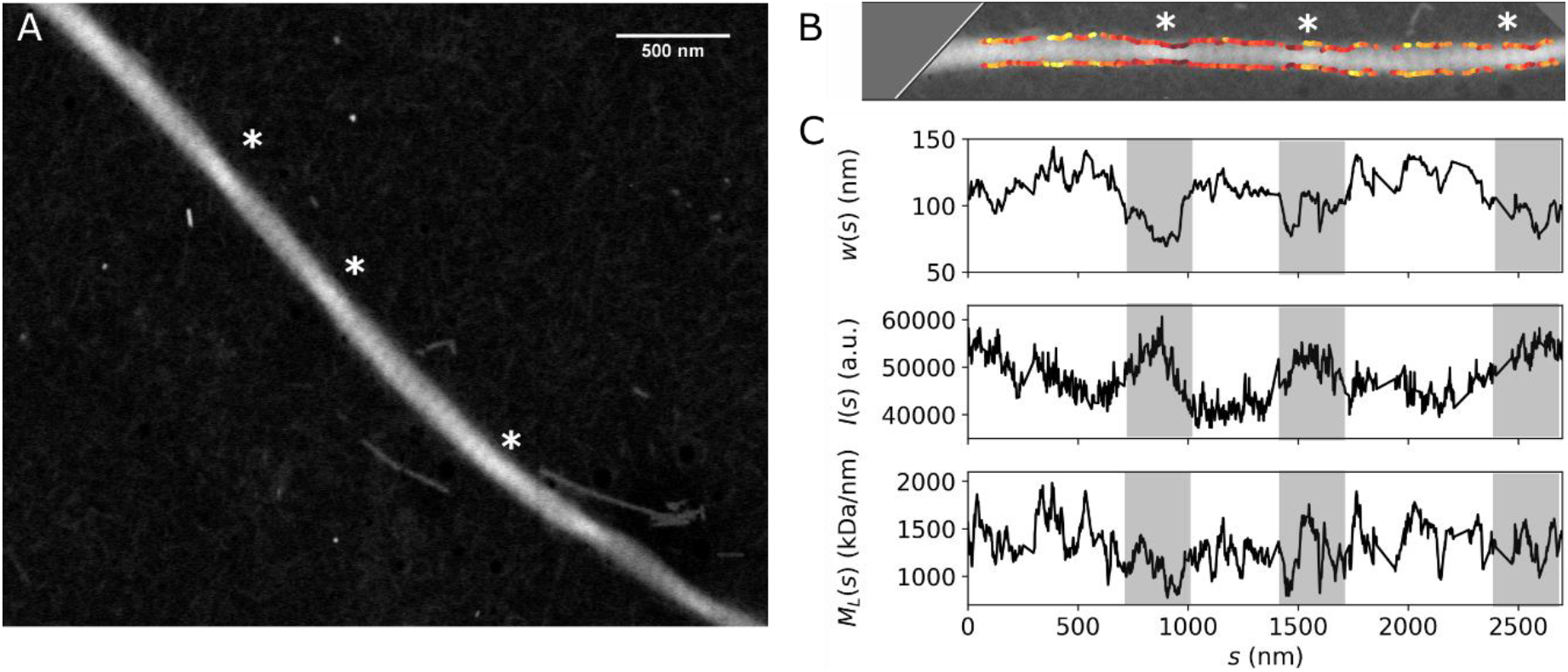
Fibril twisting. (A) STEM image of a single fibril from bovine telocollagen. The asterisks indicate the points where the intensity is enhanced and the morphology suggests the presence of fibril twists. (B) Cropped STEM image (shown in A), where the fibril has been rotated to be horizontal, and the contrast is optimized to show the twisting. The edge tracking is overlaid, with the *M*_*L*_(*s*) color coded in the same range shown in (C). (C) Axial profiles for fibril *M*_*L*_, *w*, and intensity *I*. The gray rectangles serve as a visual aid to identify the regions where fibril twisting is observed in the image shown in (B).

## CONCLUSIONS

We have developed a semi-automated algorithm to perform a detailed quantitative mass analysis of filamentous proteins from scanning transmission electron microscopy (STEM) images. Although different methods can be used for measuring the molecular packing structure of proteins, *e.g.*, small angle scattering and turbidity techniques, they rely on bulk measurements and often involve assumptions about the network or protein structure to support the data analysis.^36,37^ The spatial resolution and correlative image-mass analysis of our approach allows us to retrieve, in a label-free manner, information about the filament mass per length, molecular packing density, and about features related to the axial and longitudinal assembly of the protein. We have demonstrated the potential of this technique by studying the structure of a wide range of biopolymers important in the cellular cytoskeleton and in the extracellular matrix. Our findings demonstrate that STEM has a wide mass range, from 17 kDa/nm for Fd bacteriophages and actin filaments to 20 MDa/nm for collagen bundles. A major advantage of the automated analysis over more traditional manual analysis approaches is the ease of obtaining high statistics where maps of the protein mass can be correlated with the local filament morphology. Our analysis of collagen fibrils shows that this approach is ideal for polymorphic systems of proteins that form filaments with a hierarchical, three-dimensional packing structure. There are many examples of such hierarchical fibril-forming proteins in nature where the hierarchy is essential for the mechanical and biological function. Axially periodic packing for instance ensures toughness and biological recognition in collagen^38,39^ and high resilience in fibrin.^40^ Recently there is a growing effort in the chemistry community to design synthetic fibril-forming systems that mimic these features, based on peptides, DNA, or synthetic molecules.^41–43^ STEM imaging coupled with our automated analysis provides an ideal tool to investigate the self-assembly processes of both natural and synthetic systems to aid in the design of filamentous (bio)materials.

## METHODS

### Actin sample preparation

The assembly of actin filaments was triggered by mixing globular (G-)actin (7.5 μM) purified from rabbit skeletal muscle^44^ in an actin polymerization buffer solution containing 80 mM piperazine-N, N’-bis(2-ethanesulfonic acid) (PIPES), 4 mM MgCl_2_, 75 mM KCl, and 1 mM EGTA, at pH 6.8. Actin was allowed to assemble for 1 hour at room temperature. Before deposition, the filaments were diluted to a final concentration of 1 μM in actin polymerization buffer. A 5 μL drop of the actin solution was deposited on electron microscopy grids, allowed to adsorb for 1 min at room temperature, rinsed 5 times with milliQ water, and blotted dry.

### Microtubule sample preparation

Lyophilized porcine brain tubulin (Cytoskeleton, Denver USA) was resuspended at 50-100 μM in MRB80 buffer (80 mM PIPES, 4 mM MgCl_2_ and 1 mM EDTA, at pH 6.8), snap-frozen and stored at −80 °C. Stabilized microtubule seeds were prepared using the slowly hydrolysable GTP analogue guanylyl-(α,β)-methylene-diphosphonate (GMPCPP), following the double cycle protocol as described in Ref.^45^ and using 20 μM tubulin of which 12% was Rhodamine-labelled. To increase the average microtubule length, 20 μM tubulin was mixed with 1 μL of the GMPCPP seeds and 1.7 mM GTP in MRB80 buffer. Following incubation for 30-60 minutes at 37 °C, a 5 μL drop of the microtubule solution was deposited on electron microscopy grids, allowed to adsorb for 1 min at room temperature, rinsed 5 times with milliQ water, and blotted dry.

### Collagen sample preparation

Collagen networks were reconstituted by polymerizing commercially available collagen extracted from bovine skin, with and without telopeptides – TeloCol and FibriCol respectively – (CellSystems, Germany). To trigger polymerization, the collagen solution was mixed on ice with 0.1 M sodium hydroxide (NaOH) and with phosphate buffer saline (PBS) to reach a final pH of 7.3-7.4, as measured by a pH meter. MilliQ water was added to top up the solution to the final volume. During the pH adjustment, samples were kept on ice to prevent premature polymerization of collagen. Fibril assembly was initiated by placing the samples in a closed container (comprised of the cap of a closed Eppendorf tube placed upside down) for at least 2 hours at 37 °C. The collagen fibrils were transferred to the electron microscopy grid by peeling off the collagen gel drop surface with the grid. The sample was then rinsed 5 times with milliQ water and blotted dry.

### Scanning Transmission Electron Microscopy (STEM)

Imaging was performed using the High-Angle Annular Dark-Field mode (HAADF) on a Verios 460 electron microscope (FEI, Hillsboro, OR, USA). To achieve molecular mass determination, the collected electron beam signal was calibrated using Tobacco Mosaic Virus (TMV, kindly provided by Jean-Luc Pellequer) as an internal mass calibration standard. In all samples, a drop of 2 μL of TMV solution (25 μg/mL in PBS) was added to the carbon-coated copper grids (Ted Pella, Redding, CA, USA) prior to sample deposition. TMV was allowed to adsorb onto the grid surface for 1 minute at room temperature, rinsed 3 times with miliQ water, and blotted dry. For imaging of Fd bacteriophages (kindly provided by Pavlik Lettinga), a 5 μL drop of a solution containing Fd rods in PBS was deposited on the grid containing TMV, allowed to adsorbed for 1 min, rinsed 5 times with milliQ water and blotted dry. For the actin and microtubules samples, the grids were first pretreated by placing them in a humidified chamber for 36 hours before use, to reduce the hydrophobicity of the carbon layer, and improve sample adsorption. After TMV adsorption and washing steps, a drop of 5 μL assembly buffer was added on the grids to keep them wet until sample deposition. Grids used for collagen samples did not require these extra steps in the preparation. For all samples, the imaging was performed within 2 hours following sample preparation.

### AFM imaging

After imaging the collagen samples with STEM, the grids were imaged by Atomic Force Microscopy (AFM) in contact mode in air, using a Veeco Dimension 3100 AFM. The scanning rate was set to 0.5 Hz, with a scan resolution of 512×512 pixels. Images were acquired with a SNL cantilever with a nominal spring constant of 0.06 N/m (Bruker).

### Filament tracking and mass determination

The unprocessed HAADF images were analyzed with custom mass mapping software written in Python. The software is freely available upon request and available to download from GitHub. Images are loaded and cropped to the area of interest. Then, the edge detection and tracking is performed by selecting an appropriate size for the kernel function, which will smooth and compute the gradient of the image; the threshold for the canny edge detection algorithm is chosen with the aid of visual validation of the tracked edges. The skeleton is tracked by choosing the size of the smoothing Gaussian kernel, and the threshold for converting to the binary image that will be skeletonized is validated with the aid of visual inspection. A local linear fit is performed around each skeleton point, and the intensity profile for each cross-section is retrieved. In between each step, undesired regions can be discarded from the analysis. At this stage, results are saved, and they can be loaded into the analysis module of the software, where mass mapping can be done, or analyzed with other custom tools based on Python. For mass mapping, a final visual inspection of the quality of the tracking is performed. The calibration constant for the conversion from image intensity to mass is computed by selecting the reference structures (TMV or Fd) in the image and comparing the integrated intensity values to the known mass per length. If there are no references in the image, but the calibration constant is known, it can be manually introduced. The resulting mass per length values, intensity, as well as the width, skeleton and edge coordinates for each cross-section in the tracked filaments are saved in a results file. All of the subsequent data processing (histograms, averaging, data fitting, plotting) was performed in Python, independently from our software.

## AUTHOR INFORMATION

### Author Contributions

C.M.T and G.H.K conceived the research. C.M.T., F.B. and C.A. designed the experiments and prepared the samples for imaging. C.M.T. performed AFM and STEM imaging, analyzed data and wrote the code for the custom software. G.H.K. supervised the work and assisted in the study design and data analysis. The manuscript was written through contributions of all authors. All authors have given approval to the final version of the manuscript.

### Funding Sources

#### Notes

The authors declare no competing financial interest.

## ACKNOWLEDGMENTS

The authors thank A. Iyer, A. Szuba, V. Wollrab and A. Aufderhorst-Roberts for useful discussions; A. Lof for assistance with STEM and AFM imaging; and J-L. Pellequer and P. Lettinga for kindly gifting the TMV rods and Fd rods, respectively. This work was financially supported by the Netherlands Organization for Scientific Research (NWO) with a Topsector grant and a Veni fellowship (C. Martinez-Torres). This work was further supported by the European Research Council (Synergy grant 609822). The contribution of F. Burla and G.H. Koenderink is part of the Industrial Partnership Program Hybrid Soft Materials that is carried out under an agreement between Unilever Research and Development B.V. and the NWO.

## ABBREVIATIONS

AFM: atomic force microscopy
HAADF: high angle annular dark field
STEM: scanning transmission electron microscopy
TMV: tobacco mosaic virus

## ASSOCIATED CONTENT

### Supporting Information

Supporting figures are available free of charge *via* the Internet at http://pubs.acs.org., including: mass per length values for Fd bacteriophages, quantitative analysis of AFM imaging of collagen fibrils, and a compilation of HAADF images from (a)telocollagen. (PDF file).

A version of the code used for filament tracking and mass determination that will be maintained/updated is available at the Github repository https://github.com/cristina-mt/fias.

## Graphical Table of Contents

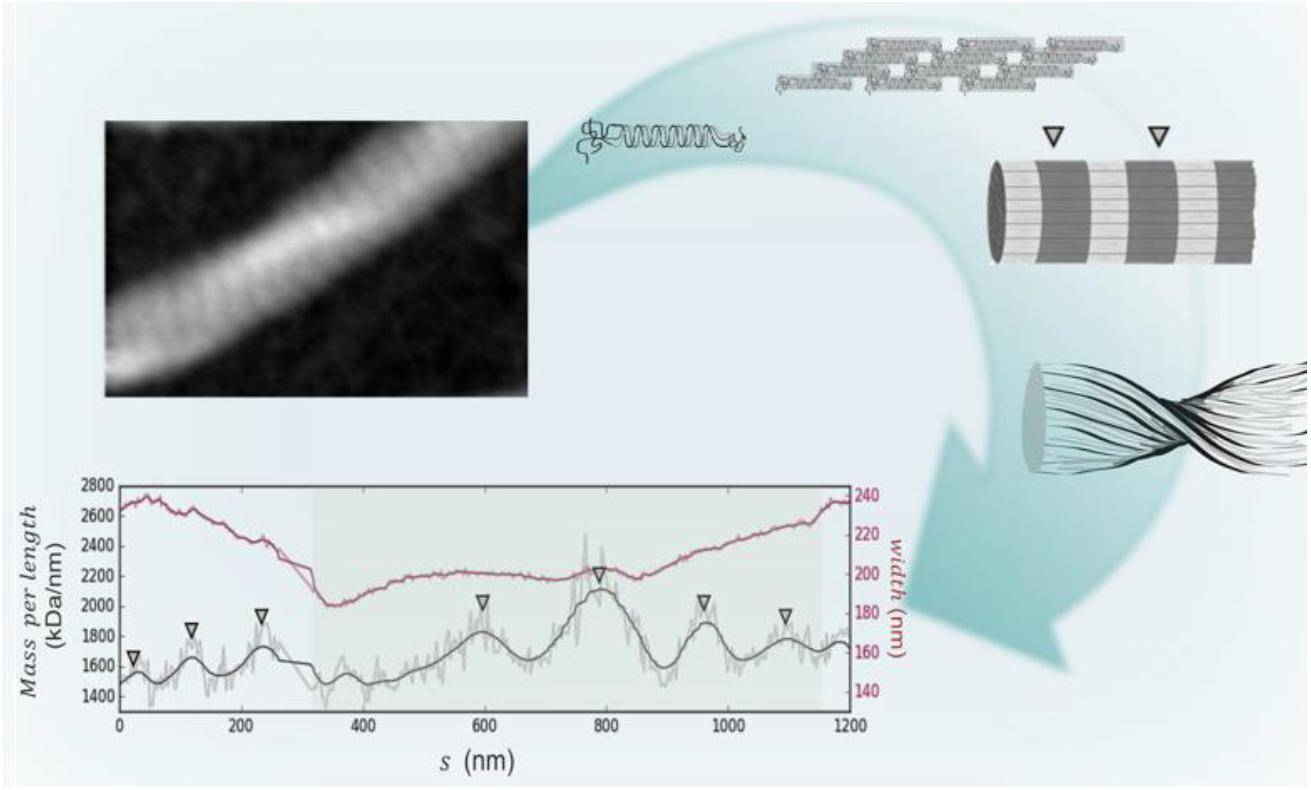

## REFERENCES

1. Buehler, M.J.; Yung, Y.C. Deformation and Failure of Protein Materials in Physiologically Extreme Conditions and Disease. Nat. Mater. 2009, 8, 175–188.

2. Schwarz, U.S.; Safran, S.A. Physics of Adherent Cells. Rev. Mod. Phys. 2013, 85, 1327–1381.

3. Burla, F.; Mulla, Y.; Vos, B.E.; Aufderhorst-Roberts, A.; Koenderink, G.H. From Mechanical Resilience to Active Material Properties in Biopolymer Networks. Nat. Rev. Phys. 2019, 1, 249–263.

4. Wall, J.S.; Hainfeld, J.F. Mass Mapping with the Scanning Transmission Electron Microscope. Ann. Rev. Biophys. Biophys. Chem. 1986, 15, 355–376.

5. Muller, S.A.; Engel, A. Structure and Mass Analysis by Scanning Transmission Electron Microscopy. Micron 2001, 32, 21–31.

6. Engel, A. Molecular Weight Determination by Scanning Transmission Electron Microscopy. Ultramicroscopy 1978, 3, 273–281

7. Muller, S.A.; Goldie, K.N.: Burki, R.; Haring, R.; Engel, A. Factors Influencing the Precision of Quantitative Scanning Transmission Electron Microscopy. Ultramicroscopy 1992, 46, 317–336.

8. Freeman, R.; Leonard, K.R. Comparative Mass Measurement of Biological Macromolecules by Scanning Transmission Electron Microscopy. Journal of Microscopy 1981, 122, 275–86.

9. Bullitt, E.S.A; DeRosier, D.J.; Coluccio, L.M.; Tilney, L.G. Three-Dimensional Reconstruction of an Actin Bundle. J. Cell Biol. 1988, 107, 597–611.

10. Nojima, D.; Linck, R.W.; Egelman, E.H. At Least One of the Protofilaments in Flagellar Microtubules Is Not Composed of Tubulin. Curr. Biol. 1995, 5, 158–167.

11. Holmes, D.F.; Graham, H.K.; Trotter, J.A.; Kadler, K.E. STEM/TEM Studies of Collagen Fibril Assembly. Micron 2001, 32, 273–285.

12. Tonino, P.; Simon, M.; Craig, R. Mass Determination of Native Smooth Muscle Myosin Filaments by Scanning Transmission Electron Microscopy. J. Mol. Biol. 2002, 318, 999–1007.

13. Goldsbury, C.; Baxa, U.; Simon, M.N.; Steven, A.C.; Engel, A.; Wall, J.S.; Aebi, U.; Muller S.A. Amyloid Structure and assembly: Insights from Scanning Transmission Electron Microscopy. J. Struct. Biol. 2011, 173, 1–13.

14. Bozec, L.; van der Heijden, G.; Horton, M. Collagen Fibrils: Nanoscale Ropes. Biophys. J. 2007, 92, 70–75.

15. Canny, J. A Computational Approach to Edge Detection. IEEE Trans. Pattern Anal. Mach. Intell. 1986, 8, 679–968.

16. Mallat, S.; Hwang, W.L. Singularity Detection and Processing with Wavelets. IEEE Trans. Inf. Theory. 1992, 38, 617–643.

17. Martinez-Torres, C.; Laperrousaz, B.; Berguiga, L.; Boyer-Provera, E.; Elezgaray, J.; Nicolini, F.E.; Maguer-Satta, V.; Arneodo, A.; Argoul, F. Deciphering the Internal Complexity of Living Cells with Quantitative Phase Microscopy: A Multiscale Approach. J. Biomed. Opt. 2015, 20, 096005.

18. Namba, K.; Stubbs, G. Structure of Tobacco Mosaic Virus at 3.6 A Resolution: Implications for Assembly. Science 1986, 231, 1401–1406

19. Zimmerman, K.; Hagedorn, H.; Heuck, C.C.; Hinrichsen, M.; Ludwig, H. The Ionic Properties of the Filamentous Bacteriophages Pf1 and Fd*. J. Biol. Chem. 1986, 261, 1653–1655.

20. Merino, F.; Pospich, S.; Funk, J.; Wagner, T.; Kullmer, F.; Amdt, H.D.; Bieling, P.; Raunser, S. Structural Transitions of F-Actin Upon ATP Hydrolysis at Near-Atomic Resolution Revealed by Cryo-EM. Nat. Struct. Mol. Biol. 2018, 25, 528–537.

21. Chou, S.Z.; Pollard, T.D. Mechanism of Actin Polymerization Revealed by Cryo-EM Structures of Actin Filaments with Three Different Bound Nucleotides. Proc. Natl. Acad. Sci. U.S.A. 2019, 13, 201807028

22. Sousa, A.A.; Leapman, R.D. Development and Application of STEM for the Biological Sciences. Ultramicroscopy. 2012, 126, 38–49.

23. Meurer-Grob, P.; Kasparian, J.; Wade, H. Microtubule Structure at Improved Resolution. Biochemistry. 2001, 40, 8000–8008.

24. Alushin, G.M.; Lander, G.C.; Kellog, E.H.; Zhang, R.; Baker, D.; Nogales, E. High-Resolution Microtubule Structures Reveal the Structural Transitions in Αβ-Tubulin Upon GTP Hydrolysis. Cell. 2014, 157, 1117–29.

25. Hoenger, A.; Thormahlen, M.; Diaz-Avalos, R.; Doerhoefer, M.; Goldie, K.N.; Muller, J.; Mandelkow, E. A New Look at the Microtubule Binding Patterns of Dimeric Kinesins. J. Mol. Biol. 2000, 297, 1087–1103.

26. Holmes, D.F.; Lu, Y.; Starborg, T.; Kadler, K.E. Collagen Fibril Assembly and Function. Curr. Top. Devel. Biol. 2018, 130, 107–142.

27. Hulmes, D.J.; Wess, T. J.; Prockop, D.J.; Fratzl, P. Radial Packing Order and Disorder in Collagen Fibrils. Biophys. J. 1995, 68, 1661–1670.

28. Zhu, J.; Kaufman, J. Collagen I Self-Assembly: Revealing the Developing Structures that Generate Turbidity. Biophys. J. 2014, 106, 1822–1831.

29. Orgel, J.; Irving, T.; Miller, A.; Wess, T. Microfibrillar Structure of Type I Collagen *In Situ*. Proc. Natl. Academ. Sci. USA. 2006, 103, 9001–9005.

30. Fratzl, P., Ed.; Collagen: Structure and Mechanics. Springer US, 2008.

31. Kadler, K.E.; Holmes, D.F.; Trotter, J.A.; Chapman, J.A. Collagen Fibril Formation. Biochem. J. 1996, 316, 1–11.

32. Fraser, R.D.; MacRae, T.P.; Miller, A. Molecular Packing in Type I Collagen Fibrils. J. Mol. Biol. 1987, 193, 115–125.

33. Cameron, S.; Kreplak, L.; Ruttenberg, A.D. Polymorphism of Stable Collagen Fibrils. Soft Matter, 2018, 14, 4772–4783

34. Brown, A.I.; Kreplak, L.; Rutenber, A.D. An Equilibrium Double-Twist Model for the Radial Structure of Collagen Fibrils. Soft Matter, 2014, 10, 8500–11.

35. Holmes, D.F.; Chapman, J.A.; Prockop, D.J., Kadler, K.E. Growing Tips of Type I Collagen Fibrils Formed *In Vitro* Are Near-Paraboloidal in Shape, Implying a Reciprocal Relationship Between Accretion and Diameter. Proc. Natl. Acad. Sci. USA, 1992, 89, 9855–9856

36. Ferri, F.; Calegari, G.R.; Molteni, M.; Cardinali, B.; Magatti, D.; Rocco, M. Size and Density of Fibers in Fibrin and Other Filamentous Networks from Turbidimetry : Beyond a Revisited Carr-Hermans Method, Accounting for Fractality and Porosity. Macromolecules 2015, 48, 5423–5432.

37. Abass, A.; Bell, J.S.; Spang, A.; Hayes, S.; Meek, K.M.; Boote, C. SAXS4COLL: an Integrated Software Tool for Analysing Fibrous Collagen-Based Tissues. J. Appl. Cryst., 2017, 50, 1235–1240.

38. Buehler, M. Nature Designs Though Collagen: Explaining the Nanostructure of Collagen Fibrils. Proc. Natl. Acad. Sci. USA, 2006, 106, 12285–12290.

39. Perimal, S.; Antipova, O.; Orgel, J. Collagen Fibril Architecture, Domain Organization, and Triple-Helical Conformation Govern Its Proteolysis. Proc. Natl. Acad. Sci. USA, 2008, 105, 2824–2829.

40. Brown, A.E.; Litvinov, R.I.; Discher, D.E.; Purohit, P.K.; Weisel, J.W. Multiscale Mechanics of Fibrin Polymer: Gel Stretching with Protein Unfolding and Loss of Water. Science, 2009, 325, 741–4

41. Kouwer, P.H.; Koepf, M.; Le Sage, V.A.; Jaspers, M.; van Buul, A.M.; Eksteen-Akeroyd, Z.H.; Woltinge, T.; Schwartz, E.; Kitto, H.J.; Hoogenboom, R.; Picken, S.J.; Nolte, R.J.; Mendes, E.; Rowan, A.E. Responsive Biomimetic Networks from Polyisocyanopeptide Hydrogels. Nature, 2013, 493, 651–5

42. Schuldt, C.; Schnauß, J.; Händler, T.; Glaser, M.; Lorenz, J.; Golde, T.; Käs, J.A.; Smith, D.M. Tuning Synthetic Semiflexible Networks by Bending Stiffness. Phys Rev. Lett., 2016, 117, 197801.

43. Knowles, T.P.; Mezzenga, R. Amyloid Fibrils as Building Blocks for Natural and Artificial Functional Materials. Adv. Mater. 2016, 28, 6546–61.

44. Alvarado, J.; Koenderink, G.H. Reconstituting Cytoskeletal Contraction Events with Biomimetic Actin-Myosin Active Gels. In Building a cell from its component parts. Ross, J., Marshall, W., Eds.; Elsevier, Methods in Cell Biology 128: Amsterdam, 2015; pp. 83–103.

45. Preciado Lopez, M.; Huber, F.; Grigoriev, I.; Steinmetz, M.O.; Akhmanova, A.; Dogterom, M.; Koenderink, G.H. *In Vitro* Reconstitution of Dynamic Microtubules Interacting with Actin Filament Networks. Methods in Enzymology, 2014, 540, 301–20.

