## Supplementary material for "Revealing the Assembly of Filamentous Proteins with Scanning Transmission Electron Microscopy": Supllemental figures

**Supporting Information**

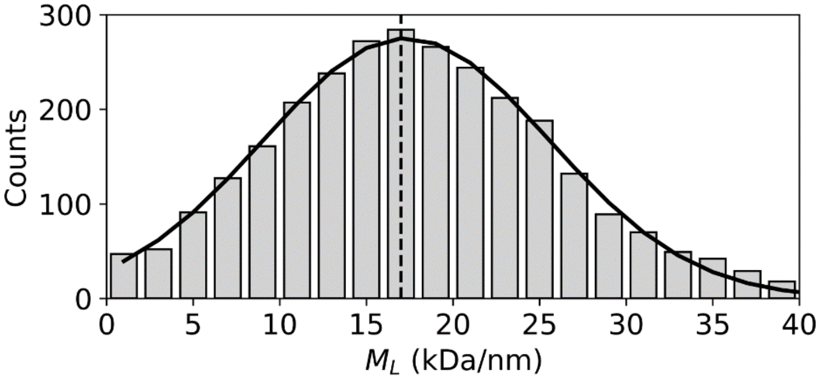


**Figure S1. Distribution of mass per length values for Fd-bacteriophage filaments, used as an internal calibration standard.** The solid black line shows a Gaussian curve fit with an average centered around $<M_L> = 17.3 \pm 8.2$ kDa/nm, and the dashed line shows the $M_L$ value (17 kDa/nm) known from the virus structure.[1]

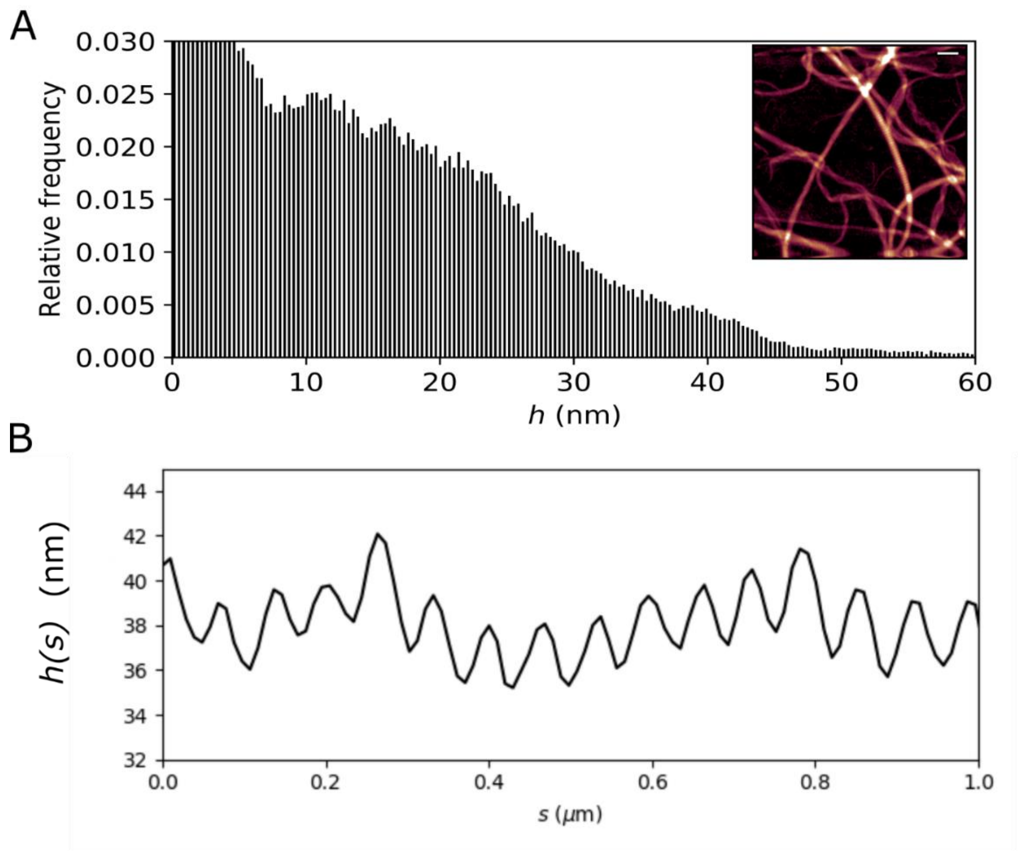


**Figure S2. Atomic force microscopy of collagen fibrils.** (A) Distribution of height *(h)* values for the AFM image of collagen fibrils shown in the inset and in Fig. 3B. There is a large majority of thin fibrils (up to 45 nm) and a few thicker ones (up to 60 nm). (B) Representative height profile for a fibril showing the characteristic D-banded pattern, where the height oscillations correspond to the D-periodicity of gap and overlap regions with an average spacing $\langle D \rangle = 65.4 \pm 5.4$ nm.

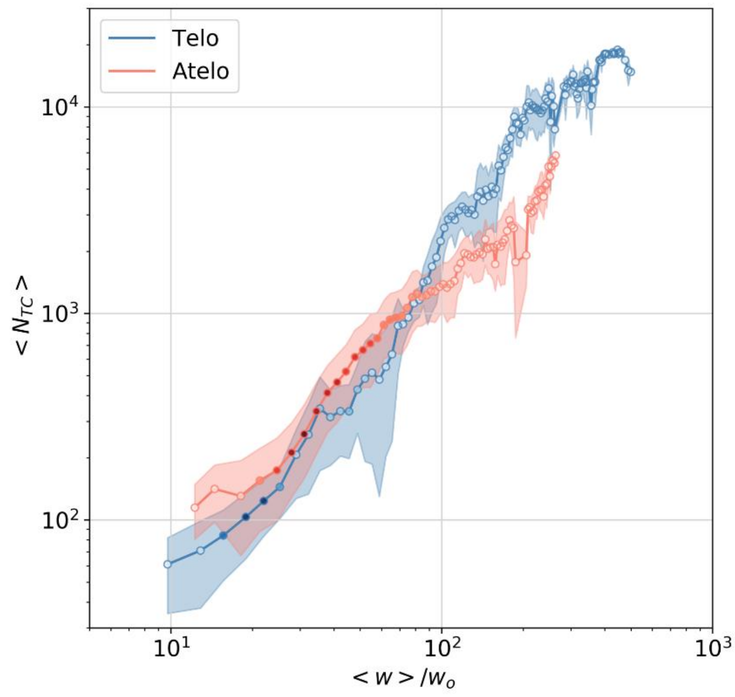


**Figure S3**. **Collagen fibril packing regimes.** Average number of tropocollagen molecules, $N_{TC}$ as a function of the normalized fibril width w/$w_o$, with $w_o$ = 1.5 nm, $m_o$ = 1.15 kDa/nm for atelo- and $m_o = 1.17$ kDa/nm for telo- collagen. The error bars represent the standard deviation. *n* = 15695 (telocollagen) and *n* = 41730 (atelocollagen) segments. Same data as in Fig. 4B in log-log representation and normalized.

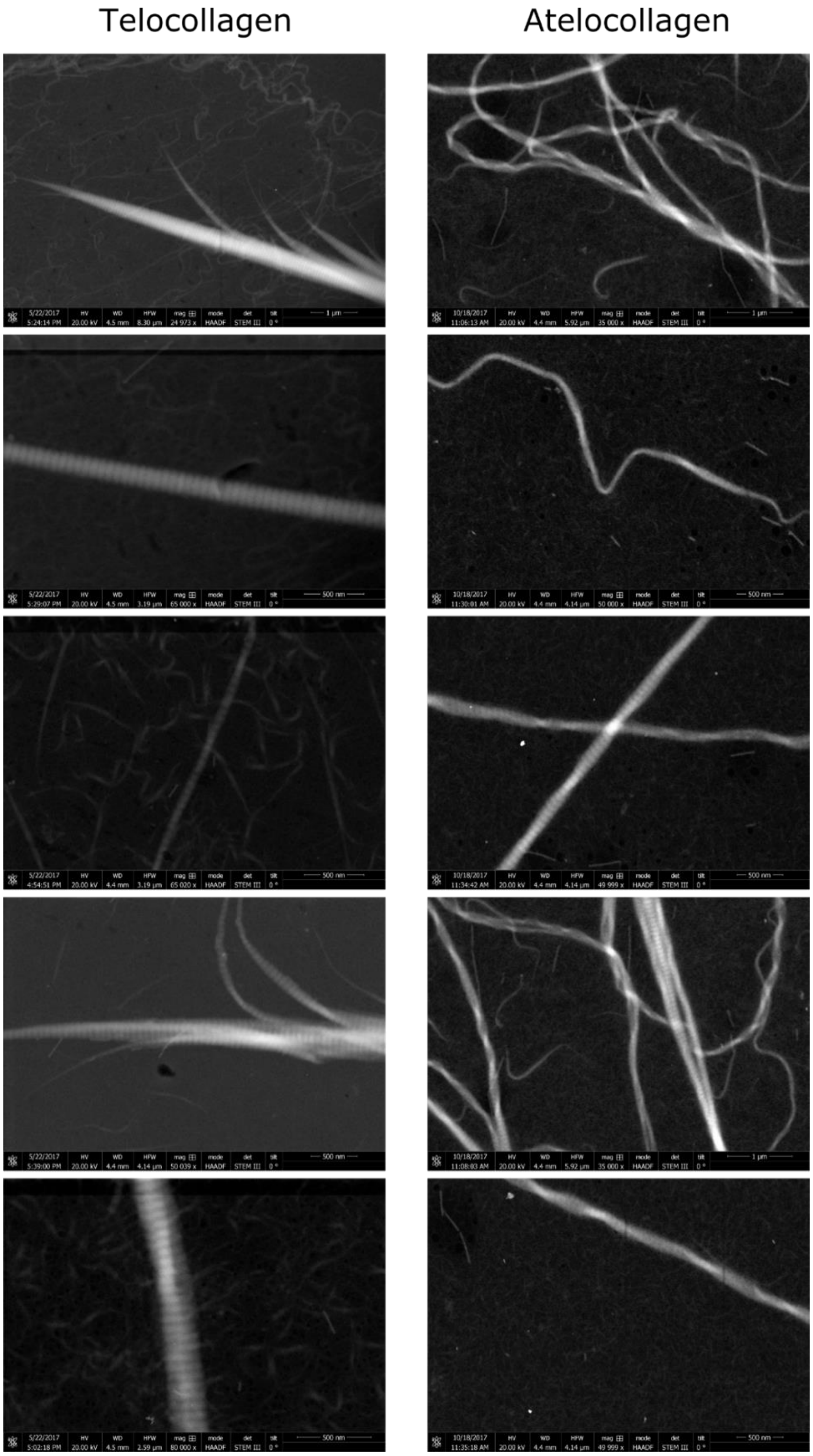


**Figure S4. Compilation of HAADF images from telocollagen (left) and atelocollagen (right) fibrils**. Each column shows 5 random images taken from the dataset of each type of collagen. Note that both collagen types form polymorphic fibrils that vary in width and $M_L$, and in the absence or presence of D-banding.

### References Supporting Information

1. Zimmerman, K.; Hagedorn, H.; Heuck, C.C.; Hinrichsen, M.; Ludwig, H. The Ionic Properties of the Filamentous Bacteriophages Pf1 and Fd*. J. Biol. Chem. 1986, 261, 1653-1655.
